# Development of a human glioblastoma model using humanized DRAG mice for immunotherapy

**DOI:** 10.1101/2023.02.15.528743

**Authors:** Rashmi Srivastava, Alireza Labani-Motlagh, Apeng Chen, Fan Yang, Danish Ansari, Sahil Patel, Honglong Ji, Scott Trasti, Meghana Dodda, Yash Patel, Han Zou, Baoli Hu, Guohua Yi

## Abstract

Glioblastoma (GBM) is the most common and lethal primary brain tumor with high mortality rates and a short median survival rate of about 15 months despite intensive multimodal treatment of maximal surgical resection, radiotherapy, and chemotherapy. Although immunotherapies have been successful in the treatment of various cancers, disappointing results from clinical trials for GBM immunotherapy represent our incomplete understanding. The development of alternative humanized mouse models with fully functional human immune cells will potentially accelerate the progress of GBM immunotherapy. In this study, we developed a humanized DRAG (NOD.Rag1KO.IL2RγcKO) mouse model, in which the human hematopoietic stem cells (HSCs) were well-engrafted and subsequently differentiated into a full lineage of immune cells. Using this humanized DRAG mouse model, GBM patient-derived tumorsphere lines were successfully engrafted to form xenografted tumors, which can recapitulate the pathological features and the immune cell composition of human GBM. Importantly, the administration of anti-human PD-1 antibodies in these DRAG mice bearing a GBM patient-derived tumorsphere line resulted in decreasing the major tumor-infiltrating immunosuppressive cell populations, including CD4^+^PD-1^+^ and CD8^+^PD-1^+^ T cells, CD11b^+^CD14^+^HLA-DR^+^ macrophages, CD11b^+^CD14^+^HLA-DR^−^CD15^−^ and CD11b^+^CD14^−^ CD15^+^ myeloid-derived suppressor cells, indicating the humanized DRAG mouse model as a useful model to test the efficacy of immune checkpoint inhibitors in GBM immunotherapy. Together, these results suggest that humanized DRAG mouse models are a reliable preclinical platform for brain cancer immunotherapy and beyond.

## Introduction

Glioblastoma (GBM), the most common and lethal primary brain tumor, is characterized by the highest mortality rates and a short median survival rate among all cancers. Immunotherapies have yielded tremendous success in many types of cancers[1; 2]. Unfortunately, the hope of using immunotherapy for brain tumor treatment has been smashed due to the recent frustrating results of clinical trials[3; 4]. This outcome discrepancy is partly due to the lack of a suitable animal model for brain cancer research. Although there are many etiological and physiopathological similarities and relevance between mouse and human brain tumors[5], the commonly used rodent models of gliomas could not faithfully recapitulate the human tumors.

Nevertheless, immunodeficient (e.g., NOD-SCID) mice have been successfully used to transplant human brain tumor cells or tissues[6; 7; 8]. These human tumors-xenografted immunodeficient mice can mainly mimic the human tumor environment, thus providing a powerful tool to study the therapeutic efficacy of the anti-tumor drugs. However, tumor progression is generally the outcome of the interplay between the tumor cells and the immune cells in clinical settings. While in these immunodeficient mice, the absence of an immune system makes testing immunotherapies for brain tumors unfeasible since the success of the immunotherapies mainly involves the modifications/activation of immune cells to improve their tumor-killing capacity. Recently, several humanized mouse models that harness the human immune system in immunodeficient mice have been developed to study specific types of cancers[9; 10; 11; 12]. In these humanized mouse models, human peripheral blood mononuclear cells (PBMCs) or hematopoietic stem cells (HSCs) have been successfully engrafted in different backgrounds of immunodeficient mice, and the engrafted/differentiated human cells demonstrated various levels of immune responses for cancer cell killing. However, the synergy between the brain tumor cells and the tumor-relevant human immune cells, such as myeloid cells and regulatory T cells development in those mice, has not been fully characterized. Therefore, developing an alternative humanized mouse model with fully functional human immune cells for studying brain tumors, particularly brain tumor immunotherapy, is of clinical significance.

In this study, we generated a new humanized mouse model by engrafting the human DR4^+^ HSCs into DRAG mice. The HSCs were well-engrafted and differentiated into an entire lineage of human immune cells, including CD4^+^ and CD8^+^ T cells, B cells, natural killer (NK) cells, Monocytes/macrophages, and dendritic cells (DCs). Based on this DRAG mouse, we then established a humanized GBM model by intracranially implanting patient-derived tumor neurospheres. Pathological and immunological characterization revealed that this humanized GBM faithfully recapitulates the hallmark features of patient tumors including the presence of a functional immune microenvironment. Notably, a pilot study of immune checkpoint blockade therapy showed this humanized GBM model as a valuable tool for understanding GBM biology and preclinically testing immunotherapy.

## Results

### Engraftment and differentiation of human immune cells in DRAG mice

To generate humanized DRAG mice, we irradiated the mice with a dose of 250 centigray (cGy) and intravenously administered 2×10^5^ CD34^+^hDR4^+^ HSCs after 6 hours of irradiation (**Fig. 1A**). The mice were monitored for more than 12 weeks, waiting for the HSCs to differentiate. After 12 weeks, we collected blood and spleen from the DRAG mice to characterize human immune cells by staining and flow cytometry. We also detected human CD45^+^ (hCD45) cell differentiation in the blood samples and spleen of the HSC-engrafted mice.

**Figure 1.**
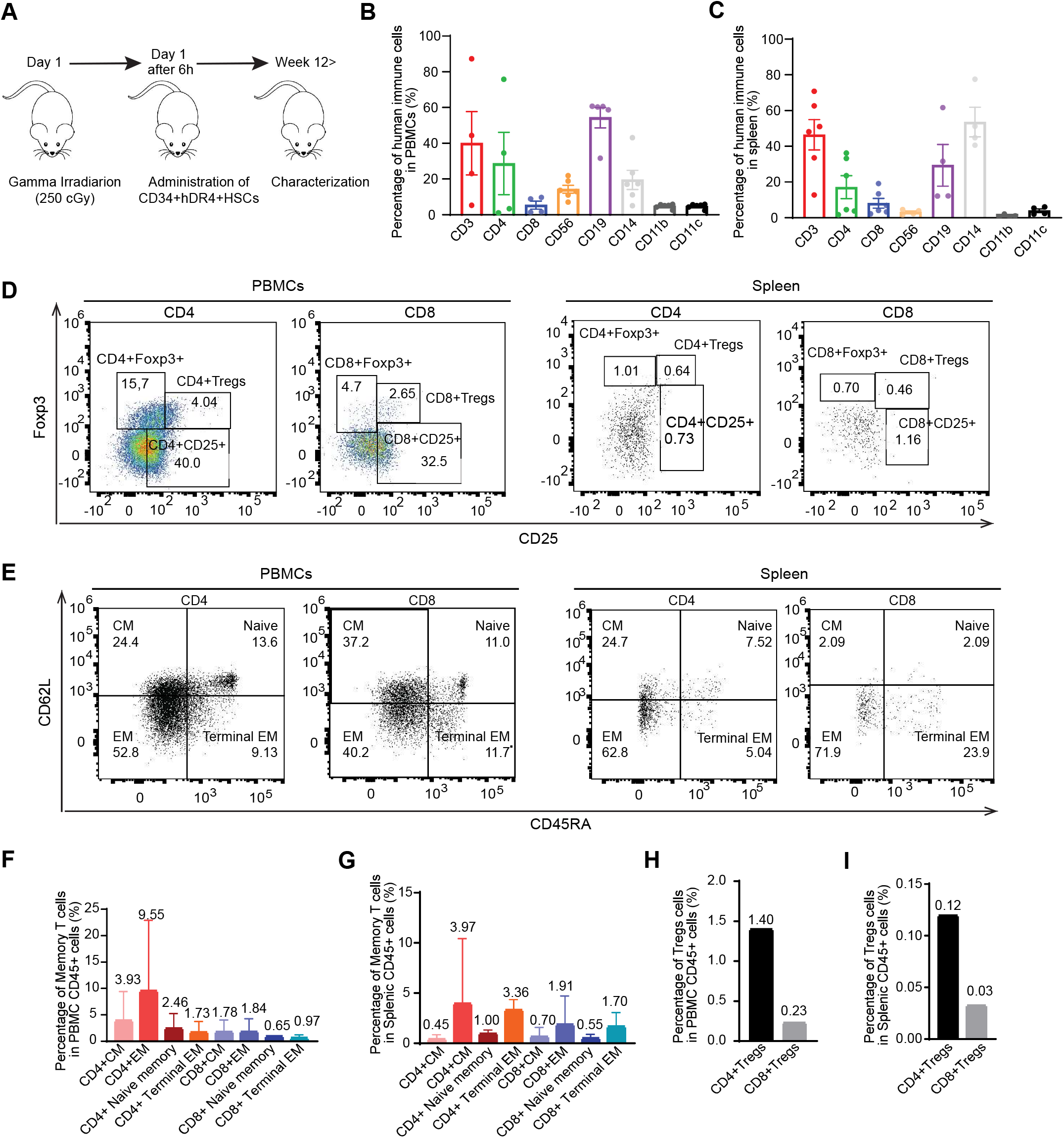
Engraftment and differentiation of human immune cells in DRAG mice. **(A)** The mice were administered with hCD34^+^hDR4^+^ hematopoietic stem cells 6 hours after gamma irradiation with a dose of 250 cGy. After 12 weeks, human immune cells in the PBMCs and spleen were characterized with flow cytometry. **(B)** Differentiation of human immune cells and their distribution in the blood of the engrafted DRAG mice. Human stem cells differentiated into CD45^+^, CD3^+^, CD4^+^, CD8^+^, CD56^+^, CD19^+^, CD14^+^, CD11b^+^ and CD11c^+^ cells with different frequency. **(C)** Differentiation of human immune cells and their distribution in the spleens of the engrafted DRAG mice. **(D)** Differentiation and the percentages of regulatory T cells (CD25^+^Foxp3^+^CD4^+^ and CD25^+^Foxp3^+^CD8^+^ Tregs) in PBMCs and in spleens. Only one mouse is shown as the representative. **(E)** Differentiation and the percentages of memory T cells (both CD4+ and CD8+) in PBMCs and in spleens. Only one mouse is shown as the representative. **(F)** Statistics of the percentages of various memory T cells in PBMCs. **(G)** Statistics of the percentages of various memory T cells in spleens. **(H)** Statistics of the percentages of Tregs in PBMCs. **(I)** Statistics of the percentages of Tregs in spleens. For all the statistical figures, each bar represents 4-6 mice.

The percentage of hCD45+ cells ranged from 0.3 to 11.6 in PBMCs (**Suppl. Fig. 1 A**) and 7.1 to 13.2 in the spleens (**Suppl. Fig. 1 A**). However, only one mouse had 57.1% and 62.2% of hCD45^+^ cells in the peripheral blood and spleen, respectively (**Suppl. Fig. 1 B and Suppl. Fig. 2 B**). This discrepancy could be due to several factors, including the individual genetic variance of the mice and the tail vein administration of human hematopoietic stem cells. We observed human lymphocytes (such as CD4^+^ T cells, CD8^+^ T cells, NK cells, and B cells), CD14^+^ cells, CD11b, and CD11c-expressing cells in the PBMCs and spleens with different frequencies (**Fig. 1B, C; Suppl. Fig. 1 and 2; Suppl. Fig. 3B and 3C** for gating strategy). The median percentage of CD3^+^ cells, CD4^+^ T cells, CD8^+^ T cells, NK cells, B cells, monocytes, CD11b^+^ cells, and CD11c^+^ cells in the PBMCs was 40, 28.6, 5.3, 14.2, 54.3, 19.5, 4.7, and 4.7, respectively (**Fig. 1B, C; Suppl. Fig. 1 C-H**). The median percentage of these cells in the spleens was 46.4, 17.1, 8.1, 3.1, 29.3, 53.5, 0.9, and 3.9, respectively (**Fig. 1B, C; Suppl. Fig. 2 B-G**). Moreover, we could detect human Foxp3^+^CD25^+^ regulatory CD4^+^ and CD8^+^ T cells (Tregs) in both tissues with a very low percentage. The percentage of regulatory CD4^+^ and CD8^+^ T cells was 1.4 and 0.23 in the PBMCs and 0.12 and 0.033 in the spleen (**Fig. 1 D, H, I and Suppl. Fig. 3A** for gating strategy). Furthermore, we examined human memory CD4^+^ and CD8^+^ T cells in both PBMCs and spleen of the mice (**Fig. 1 E, F, and G**), including CD62L^+^CD45RA^−^ (central memory), CD62L^+^CD45RA^+^ (naïve memory), CD62L^−^CD45RA^−^ (effector memory) and CD62L^−^CD45RA^+^ (terminal effector memory) cells. In the PBMCs, CD4^+^ central and effector memory cells were the highest in the PBMCs; however, CD4^+^ effector and terminal effector cells in the spleen presented the highest memory cell subsets in the engrafted DRAG mice (**Fig. 1 E, F, G, and Suppl. Fig. 3A** for gating strategy). These results suggest that the DRAG mice developed a full lineage of immune cells as in humans.

**Figure 2.**
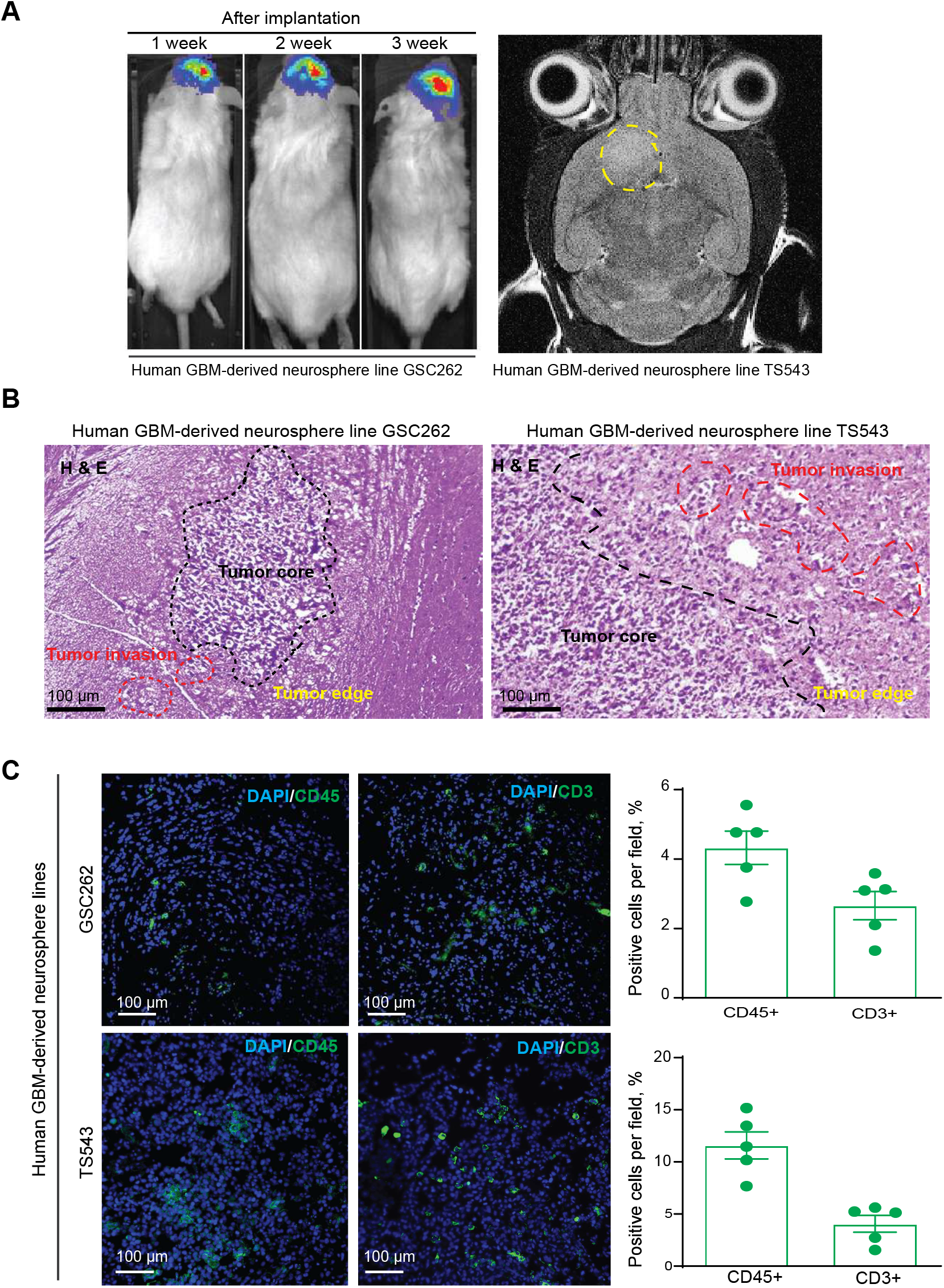
Establishment of xenograft tumors using DRAG mice bearing GBM patient-derived tumor cells. **(A)** The luminescence images show a DRAG mouse bearing TS543 cells after implantation at three different time points; MRI from a DRAG mouse after intracranial implantation of GSC262 cells. T2 sequences demonstrate infiltrative tumors in the mouse brain (yellow line). **(B)** H&E analyses of tumor sections show the core, edge, and invasion regions of the tumors derived from the DRAG mice bearing TS543 and GSC262 cells. **(C)** Representative immunofluorescence staining images for CD45^+^ and CD3^+^ cells in tumor sections from the DRAG mice bearing TS543 and GSC262. Quantitation of indicated cells in tumor regions and each dot represents 1 field of the tumor regions from these tumors.

**Figure 3.**
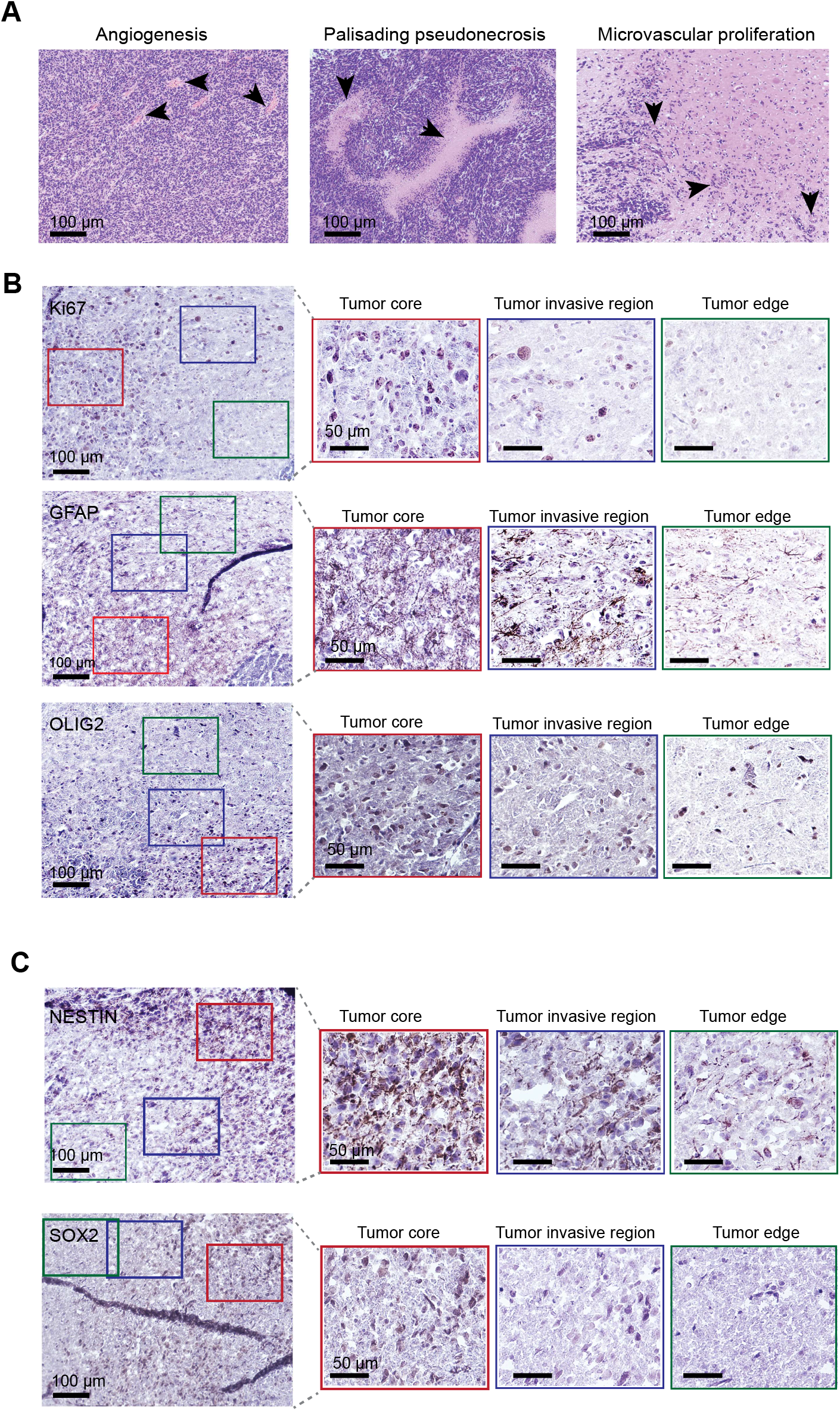
Histopathological analysis of human GBM from humanized DRAG mice. **(A)** Representative H&E images harvested from humanized DRAG mice showing microvascular proliferation (MVP) in the PVN niche, pseudopalisading necrosis in the hypoxic niche, and angiogenesis and aggressive MVP in the invasive region. **(B, C)** Representative images of immunohistochemical staining of normal brain region (green box), tumor core region (red box), and invasive tumor region (blue box) showing expressions of high proliferative index Ki67, astrocyte and oligodendrocyte lineage marker genes GFAP and Olig2, neural stem cell lineage markers Nestin and SOX-2 expression in GBM tumors.

### Humanized DRAG mice support the growth of GBM patient-derived tumor neurospheres

To elucidate the function of the differentiated human immune cells in such engrafted DRAG mice, we tested whether these immune cells can respond to patient GBM cells that were intracranially implanted in the brains of the DRAG mice. First, two GBM patient-derived neurosphere lines, GSC262 and TS543, were intracranially implanted in the DRAG mouse brains respectively. Tumor growth was monitored using the bioluminescence in vivo imaging system (IVIS) and magnetic resonance imaging (MRI) in these mice (**Fig. 2A**). Second, hematoxylin and eosin (H&E) staining showed spatial heterogeneity including tumor cores with a high density of malignant cells, tumor edges, and invasive areas where tumor cells infiltrated the normal brain parenchyma (**Fig. 2B**). To examine tumor-infiltrating immune cells that were differentiated from CD34^+^hDR4^+^ HSCs, we performed immunofluorescence staining analysis of CD45^+^ and CD3^+^ cells in the tumor tissues from the DRAG mice bearing GSC262- and TS543-derived GBM. Of note, CD45^+^ immune cells represented 4.3 ± 1.1% and 11.6 ± 2.9% of the whole tumor tissue cellularity in GSC262- and TS543-derived GBM, respectively (**Fig. 2C**). Furthermore, CD3^+^ T cells represented 2.7 ± 0.9% and 4.0 ± 1.8% of the whole tumor tissue cellularity in GSC262- and TS543-derived GBM, respectively **(Fig. 2C)**. The percentages of these immune cells are close to the range of CD45^+^ and CD3^+^ cells in patient GBM samples [13], indicating these glioma models using DRAG mice recapitulate the human GBM immune microenvironment.

### Humanized DRAG mouse-derived tumors retaining histopathological properties of patient GBM

To assess whether the tumors derived from the humanized DRAG mice recapitulate histopathological features of patients’ GBM, H&E analysis revealed that the brain tumors displayed typical GBM pathological features – pseuodopalisading necrosis and microvascular proliferation, which are a few of the hallmark characteristics of hypoxic and perivascular niches, including neoangiogenesis in the invasive niche of the GBM tumor (**Fig. 3A**). Furthermore, histopathological characterization of these tumors documented other classical GBM properties, including a high proliferative index (Ki67) and robust expression of astrocyte and oligodendrocyte lineage marker genes, glial fibrillary acidic protein (GFAP) and Oligodendrocyte Transcription Factor 2 (Olig2) (**Fig. 3B**). The hallmark of GBM is the existence of glioblastoma stem cells (GSCs), which are able to be characterized by highly expressed neural stem cell lineage markers (such as Nestin and SOX2)[14]. Therefore, we examined these tumors and observed high expression of these genes in the tumor core (CT), the leading edge of the tumors (LE), and the invasive regions (**Fig. 3B, C**). These results indicate that the novel humanized DRAG mice model supports the GBM tumor engraftment and progression, and faithfully recapitulates the pathological and molecular features of GBM.

### High immunosuppression in the humanized DRAG mouse-derived GBM

GBM is generally considered highly immunosuppressive and resistant to immunotherapy, mainly because of a relatively low mutational burden, high percentages of M2-like protumor macrophages, and low numbers of tumor-infiltrating lymphocytes and other immune effector cells[3; 15; 16]. To examine the features of immunosuppression in the humanized DRAG mouse-derived GBM, we first analyzed infiltrating tumor-associated macrophages in tumors by flow cytometry and found 16% of CD11b^+^/HLA-DR^+^/CD14^+^ cells in the CD45^+^ cell population (**Fig 4A, B**). Notably, M2-like macrophages (CD45^+^CD206^+^) accounted for 60.8% of the CD45^+^ cells in the tumors (**Fig 4A, B**). As evident in human GBM [17], we have also observed that CD45^+^CD206^+^ M2 macrophage cells as the most predominant as compared to other myeloid cells within TILs (**Fig. 4B**). Besides these myeloid cells, we have found a significantly higher frequency of CD11b^+^/CD14^+^/HLA-DR^+^ tumor-associated macrophages (TAMs) than myeloid-derived suppressor cells (MDSCs) within the CD45^+^ immune cells (**Fig. 4B**).

**Figure 4.**
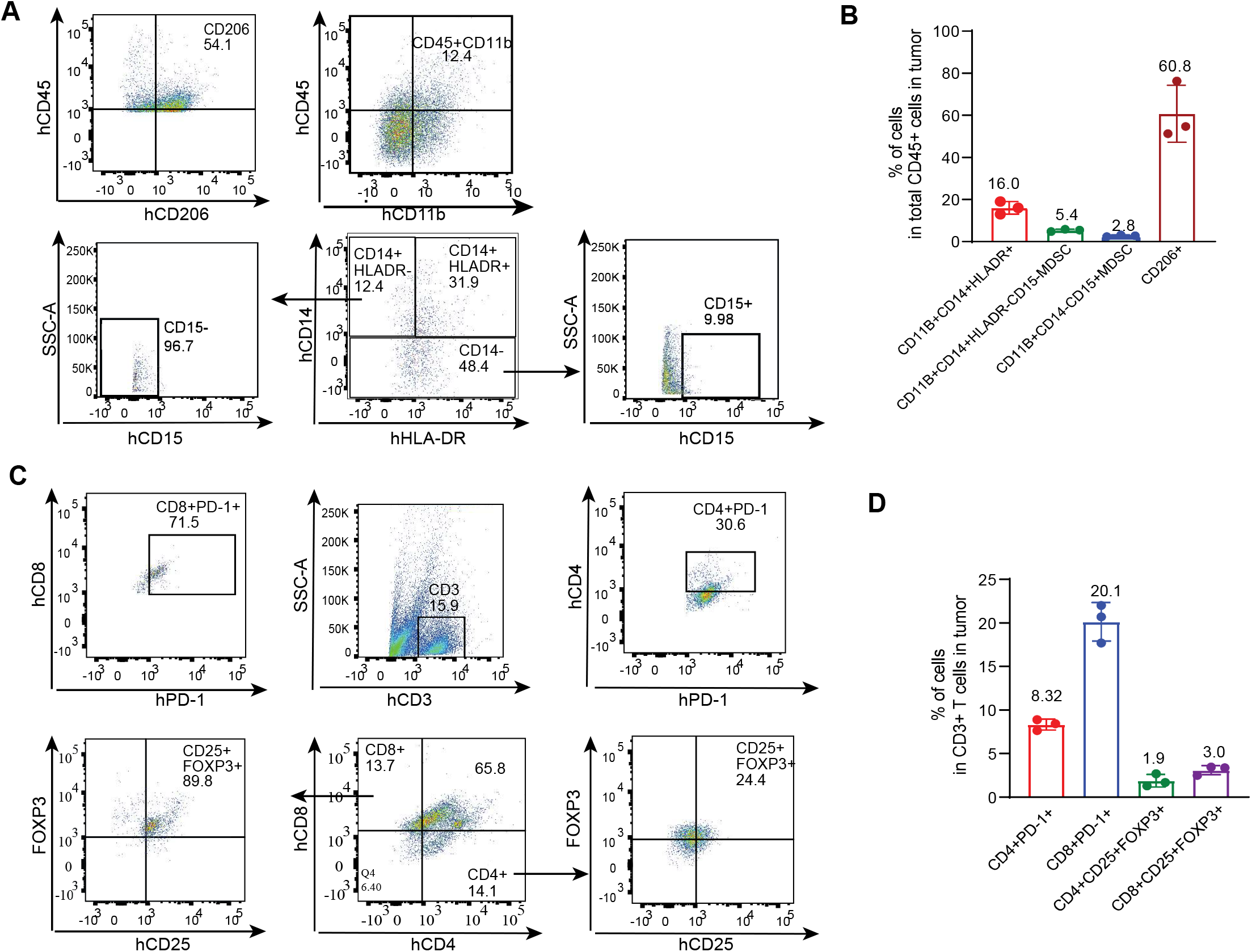
Infiltrating human lymphocytes (TILs) in the tumor derived from humanized DRAG mouse analyzed by flow cytometry. **(A)** The flow cytometry charts show the gating and percentages of the macrophage populations, including hCD45^+^/hCD206^+^, hCD45^+^/hCD11b^+^, hCD45^+^/hCD11b^+^/hCD14^+^/hHLA-DR^−^/hCD15^−^, hCD45^+^/hCD11b^+^/hCD14^+^/hHLA-DR^+^, and hCD45^+^/hCD11b^+^/hCD14^−^/hCD15^+^ in the total infiltrated cells from the tumor-derived from humanized DRAG mice. **(B)** Histograph showing the percentages of CD11b^+^/CD14^+^/HLA-DR^+^, CD11b^+^/CD14^+^/HLA-DR^−^/CD15^−^, CD11b^+^CD14^−^CD15^+^, and CD206^+^ in the total human CD45^+^ cells in the tumors. Each dot represents one mouse **(C)** The flow cytometry charts show the gating and percentages of the macrophage populations, including hCD4^+^hPD-1^+^, hCD8^+^hPD-1^+^, and hCD4^+^hCD25^+^hFOXP3^+^ cells in the hCD3^+^ immune cells infiltrated from the DRAG mice derived tumors. **(D)** Histograph showing the percentages of CD4^+^/PD-1^+^, CD8^+^/PD-1^+^, CD4^+^/CD25^+^/FOXP3^+^ and CD8^+^/ CD25^+^/FOXP3^+^ in the total human CD3^+^ cells in the TILs derived from DRAG mice bearing TS543 cells. Each dot represents one mouse.

Given that the regulatory T cells (Tregs) play a critical role in inhibiting the activation and differentiation of CD4 helper T cells and CD8^+^ cytotoxic T cells to interact against tumor-expressed antigens [18], and PD-1 is identified as a typical T cell immunosuppressive marker[19], we quantified PD-1^+^ T cells and CD25^+^FOXP3^+^ regulatory T cells among the CD3^+^T cells in the tumor immune microenvironment (TIME). Regarding CD3^+^ T cells, we have found the percentage of CD8^+^PD-1^+^ T cells is predominantly higher than that of CD4^+^PD-1^+^ T cells (**Fig. 4C, D**). We also detected the presence of CD25^+^FOXP3^+^ regulatory T cells (Tregs) in the TIME, where the frequency of CD8^+^ Tregs is higher than the CD4^+^ Tregs, while it is not significant (**Fig. 4C, D**). Taken together, these results demonstrated that GBM tumors engrafted in humanized DRAG mice not only mimic the human GBM tumor in terms of pathological and molecular features but also recapitulate the TILs for its progression.

### Anti-tumor effect of the anti-PD-1 antibody

To determine whether the human immune cells differentiated in the humanized DRAG mice have the same functions as in humans, we have compared the tumor-infiltrated immune profile in these mice injected with an anti-PD-1 antibody or an isotype IgG as control. There are no statistically significant differences between both groups regarding tumor volume and survival possibly due to the mouse number (data not shown). However, we found significant decreases in the frequency of CD4^+^PD-1^+^ and CD8^+^PD-1^+^ T cells in the anti-PD-1-treated group (**Fig. 5A, B**) when we examined the status of immune cells within the TIME by flow cytometry analysis. Furthermore, we noted that Tregs tend to decrease in number upon the treatment with the anti-PD-1 antibody (**Fig. 5C**).

**Figure 5.**
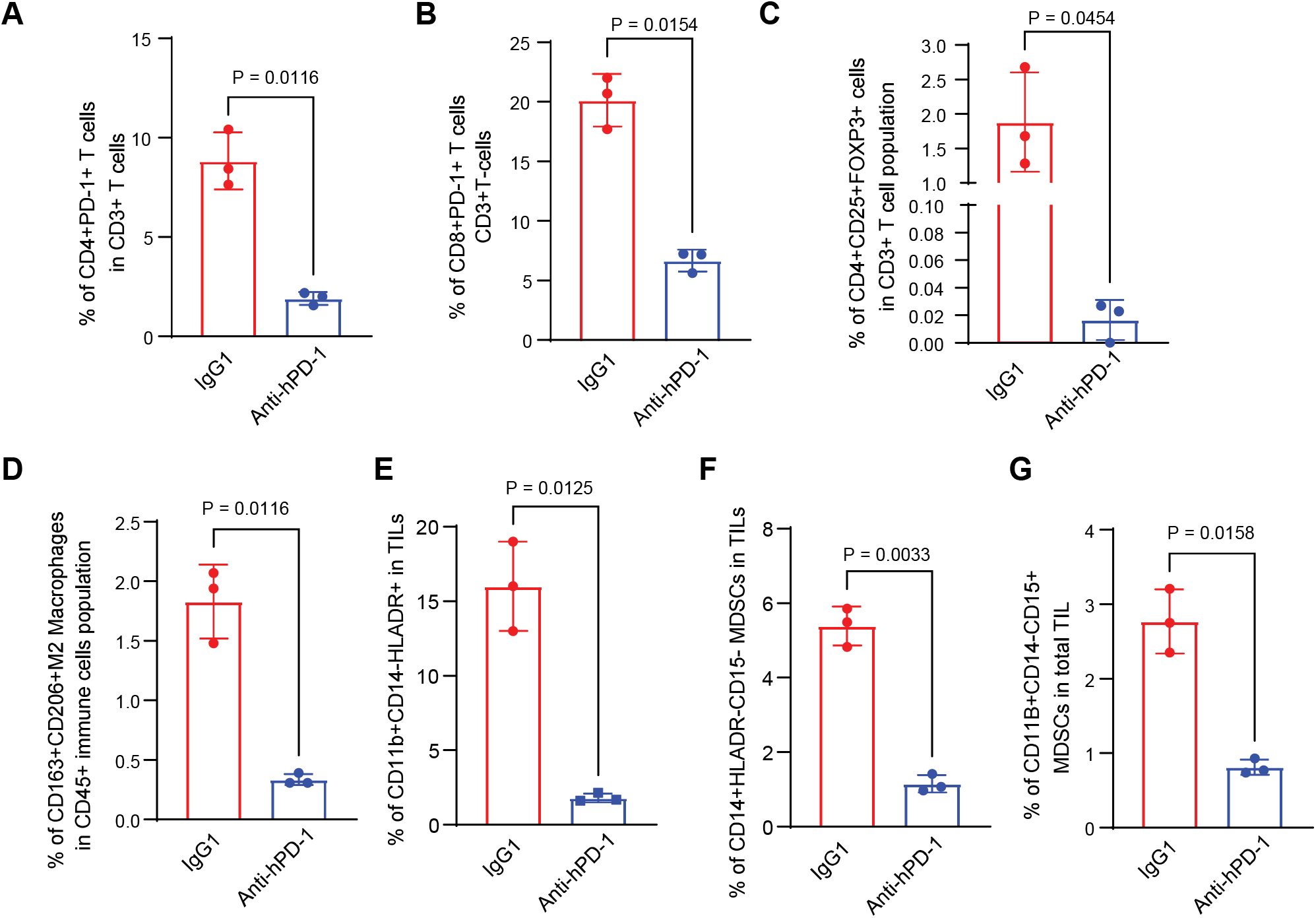
Effect of anti-PD-1 on immune cells infiltrating GBM tumors derived from DRAG mice. The Anti-PD-1 antibody and isotype IgG control salines were injected intraperitoneally at 10 mg/kg on day 14 and day 21 post-intracranially implantation of orthotopic xenografting of TS543 cells in the brain. Each dot represents one mouse. **(A)** the percentage of tumor-infiltration of human CD4^+^/PD-1^+^ cells, **(B)** CD8^+^/PD-1^+^ cells, and **(C)** CD4^+^/CD25^+^/FOXP3^+^ regulatory T cells in the human CD3^+^ cells, or **(D)** CD163^+^/CD206^+^ macrophages, **(E)** CD11b^+^/CD14^+^/HLA-DR^+^ macrophages, **(F)** CD11b^+^/CD14^+^/HLA-DR^−^/CD15^−^ and **(G)** CD11b^+^/CD14^−^/CD15^+^ MDSCs from the TILs that were isolated from the DRAG mice bearing TS543 cells with the treatment of anti-PD-1 antibody vs the isotype IgG. Each dot represents one mouse.

To test whether the treatment of anti-PD-1 antibody has affected TAM and MDSC infiltration in the tumors, we stained the cells with human TAM and MDSCs markers for flow cytometry analysis. There is no difference in the frequency of human CD45^+^ immune cells between the anti-PD-1 Ab-treated and control-IgG-treated groups, but a significantly lower frequency of CD45^+^CD163^+^CD206^+^ M2 macrophages has been observed in the anti-PD-1 Ab-treated mice (**Fig. 5D**). We also observed that the frequencies of CD11b^+^CD14^+^HLA-DR^+^ macrophages, CD11b^+^CD14^+^HLA-DR^−^CD15^−^ and CD11b^+^CD14^−^CD15^+^ MDSCs are significantly depleted upon the treatment of anti-PD-1 Ab as compared to the control-IgG Ab (**Fig. 5E-G**). These results indicate that our humanized DRAG mouse model can be used to predict the efficacy of anti-PD-1 antibodies prior to clinical treatment.

## Discussion

A good humanized mouse model that can recapitulate the clinical setting of the human cancer microenvironment is crucial not only for tumor biology research but also for the development of therapeutic strategies and agents. In this study, we established the humanized DRAG mouse model and found that the human stem cells can be well-engrafted and differentiated into a full lineage of human immune cells in these mice. Using this humanized DRAG model, we found that GBM patient-derived tumorsphere cells can successfully be engrafted and form tumors, which recapitulate the pathological features and the immune cell composition of human GBM. Importantly, we also observed this human GBM mouse model shows the response to an anti-human PD-1 antibody treatment, suggesting the feasibility of testing the efficacy of immune checkpoint inhibitors in GBM immunotherapy.

We have previously established a humanized DRAG mouse model to test a DNA vaccine protecting human cells against Zika virus infection [20]. Compared with other commonly used humanized models, such as Hu-PBL and BLT mouse models, the DRAG mouse model has several advantages: 1) constitutive expression of HLA-DR4 molecules favors engraftment of human pro-T cells in the mouse thymus for further development into mature T cells [21]; 2) immune response in DRAG mice is much more robust; 3) unlike other humanized mouse models, the Ab class switching from IgM to IgG occurs in humanized DRAG mice, which results in a more robust antibody response [22]; 4) the DRAG mice have well-developed Peyer’s patch and gut-associated lymph tissue and robustly reconstitute human T cells in the gut and other mucosal sites[23], which contains the major population of T cells. After twelve weeks HSCs injection, the mice developed a full complement of human immune cell types, including CD4^+^ and CD8^+^ T cells, B cells, and myeloid cells (**Fig. 1**). These characteristics of DRAG mice make them a valuable model for investigating how the functional human immune cells regulate cancer progression upon immunotherapies for brain tumor treatments.

Despite various types of GBM models, like syngeneic models, genetically engineered mouse models, and xenograft models, unfortunately, none of them can faithfully recapitulate the human GBM and its TIME, enabling the investigation of the efficacy of immune-checkpoints antibody inhibitors [24]. Our DRAG mice GBM model provides a promising platform to study pathological and molecular changes occurring in a GBM tumor during tumor progression. In this study, we demonstrated that the orthotopic xenograft model of GBM patient-derived tumorsphere cells intracranially implanted GBM tumor displays specific niches like hypoxic, perivascular (PVNs), GSC, and invasive niches [25]. In the PVNs, the blood vessels have evolved into microvascular proliferation (GMP), a common hallmark in GBM within the GBM-PVN. Another hallmark characteristic of GBM is pseudopalisading necrosis, within the hypoxic niche, which has also been displayed by the GBM tumor implanted in these DRAG mice.

Besides these pathological features, we also shown that the expressions of GSC markers like SOX-2, Olig-2, and Ki67 are higher in the tumor region than in the normal brain region in these GBM DRAG mice. The self-renewal pluripotency of GSC is one of its unique characteristics that are responsible for GBM heterogeneity and aggressiveness [26]. Our immunohistochemistry results have also detected the expression of GFAP and Nestin in the tumor core and invasive region, which supports that these GBM tumors display the glial- and neuronal -phenotypes for tumor progression [27; 28]. Our results suggest that this GBM tumor exhibits the pathological and histological features of intra-tumoral heterogeneity at the cellular and molecular levels for the tumor aggressiveness and progression, thus the GBM DRAG mouse model replicates these key features of a human GBM tumor [24].

This humanized DRAG mouse model is a novel model that is not limited to displaying the hallmark features of human GBM but also faithfully recapitulates the tumor-infiltrated microenvironment. Several mouse models are available, but the humanized DRAG mouse model has shown more competence in the human immune system than others [29; 30; 31]. Our model showed the presence of hCD45^+^ immune cells, including CD3^+^, CD4^+^, and CD8^+^ T cells, which are significantly associated with the progression of human GBM tumors [32]. The TIME in this mouse model also comprises a typical immunosuppressive environment by infiltrating more TAMs, MDSCs, PD-1^+^ T cells, and regulatory T cells.

In this study, we observed that anti-PD-1 antibody treatment on the GBM tumor-bearing mice significantly decreases the number of immunosuppressive cells within the TIME, indicating that this novel humanized mouse model can be used for predicting the therapeutic outcomes of immunotherapy in preclinical studies. By using this humanized mouse model platform, we can evaluate immune-checkpoint blockade/antibody therapy for human GBM treatment. Meanwhile, using this humanized GBM-DRAG mouse model, we can study not only the pathological, histological, and immune microenvironment changes caused by the tumors but also the effect of anti-PD-1 antibodies on tumor progression and TILs. Taken together, our study shows that the novel GBM DRAG mouse model can faithfully recapitulate the human GBM and represent a valuable resource for testing antitumor drugs in GBM therapy.

## Materials and Methods

### Generation of humanized DRAG mice

DRAG mice are NRG mice (or NOD.Rag1KO.IL2RγcKO mice) transgenic for human DR4 (RRID: IMSR_JAX:017914), which were purchased from Jackson Laboratories and bred in the vivarium of the University of Texas Health Science Center at Tyler. For humanization, the DRAG mice were irradiated with a dose of 250 cGy. After six hours of irradiation, 2×10^5^ hDR4+ HSCs (purchased from Stemcell Technologies Inc, and then DR4 genotyping was performed) were intravenously administrated into the mice. After 10–12 weeks, the human cell reconstitution was characterized by immune staining using a panel of biomarkers shown below.

### Isolation of PBMCs and splenic cells from DRAG mice

PBMCs were isolated from mice blood using Ficoll-Paque Plus (GE Healthcare Life Science, Piscataway, NJ, USA) procedure. After washing, the cells were treated with ACK lysing buffer (Thermo Fisher Scientific, Waltham, MA, USA) to lyse red blood cells. The spleens were ground and passed through a cell strainer. After centrifugation at 400 × g for 10 minutes, the cell pellets were subjected to ACK lysing buffer. The cells were then washed and counted before staining for flow cytometry.

### Intracranial xenograft tumor models

The intracranial xenograft tumor models were established as previously described [14; 16]. Briefly, The DRAG mice were anesthetized by intraperitoneal injection with ketamine/xylazine solution (200 mg ketamine and 20 mg xylazine in 17 mL of saline) at a dosage of 0.15 mg/10 g body weight. Then, these anesthetized mice were placed into stereotactic apparatus equipped with a z axis (Stoelting) to make a small hole in the skull 0.5 mm anterior and 3.0 mm lateral to the bregma using a dental drill. GBM patient-derived tumorsphere cells, TS543 and GSC262, diluted in 5 μL DPBS at 2 × 10^5^ cells, were intracranially injected in these DRAG mice. Animals were followed daily for the development of tumors by behavior monitoring, magnetic resonance imaging (MRI), and bioluminescent imaging.

### MRI and bioluminescent imaging

MRI and bioluminescent imaging of mice were performed at Rangos Research Center Animal Imaging Core by using the protocol as previously described [16]. Tumor appearance in the mouse brain was detected by either MRI or bioluminescent imaging. The tumor size was analyzed with ITK-SNAP. For bioluminescent imaging, DRAG mice were intraperitoneally injected with d-luciferin (150 mg/Kg; GoldBio), and images were captured by the IVIS Lumina S5 system (PerkinElmer). All animal experiments were performed with the approval of the University of Pittsburgh’s Institutional Animal Care and Use Committee (IACUC).

### Antibodies and flow cytometry

Before staining, human and mouse FC receptors were blocked using Human TruStain FcX (BioLegend, San Diego, CA, USA) and purified anti-mouse CD16/32 antibody (BioLegend), respectively. The cells were then stained with the following fluorescent-conjugated antibodies: anti-human BV421-CD45 (clone 2D1), PE/Cy7-CD45 (2D1), PE-CD3 (SK7), Alexa flour 488-CD4 (SK3), PE/Cy7-CD8 (SK1), BV605-CD8 (SK1), PE/Cy5-CD56 (HCD56), BV510-CD14 (MSE2), APC-CD11b (ICRF44), BV510-CD3 (OKT3), FITC-CD4 (A161A1), PE/Dazzle 594-CD62L (DREG-56), APC-CD45RA (HI100), BV711-CD25 (BC96), BV421-Foxp3 (206D), anti-mouse Alexa flour 700-CD45 (13/2.3), anti-mouse APC/Cy7-CD45 (30-F11) all from BioLegend; AmCyan-CD19 (SJ25C1) and PE/CF594-CD11c (BU15) from BD Bioscience (San Jose, CA, USA). The fluorescent-conjugated antibodies utilized as isotype control included mouse PE, PE/Cy7, PE/Cy5, Alexa flour 488, PE-CF594, APC, BV510, FITC, BV605, PE/Dazzle 594, BV711, BV421 conjugated IgG1(MOPC-21), mouse BV510-IgG2a (MOPC-173), mouse APC-IgG2b (MOPC-21) from BioLegend and mouse AmCyan-IgG1 (X40) from BD Bioscience. The stained cells were collected by Attune NXT (Thermo Fisher Scientific), and the data were analyzed with FlowJo (Tree Star, Ashland, OR, USA). Dead cells were removed by both forward and side scatter gating. For tumor immune cell analyses, after blocking Fc receptors, cells were stained with the following fluorescent-conjugated antibodies: anti-human PE-CD45 (2D1), FITC-CD3 (UCHT1), BV421-CD4 (RPA-T4), PE/Dazzle-594-CD8 (SK1), PerCP-PD-1 (EH12.2H7), Alexa fluor 700-CD25 (BC96), PE-FOXP3 (H1.2F3), BV421-HLA-DR (L243), APC/Fire810-CD11b (M1/70), APC-CD163 (GH1/61), PE-Cyanin7-CD206 (15-2), PE/DAZZLE-594-CD14 (HCD14) and BV711-CD15(W6D3).

### Immunohistochemistry staining

For IHC staining, brain tissues were fixed in 4% paraformaldehyde overnight and soaked in 20% sucrose solution. After 24 hours, the tumor tissue was embedded in OCT on dry ice. Brain sections were rehydrated before antigen retrieval. After quenching the endogenous peroxidase, brain sections were blocked for one hour at RT and incubated with indicated primary antibodies overnight at 4°C. Later, the brain sections were washed with blocking buffer and incubated with the respective anti-mouse/ rabbit -horseradish peroxidase (HRP)-conjugated polymer for 40 min and then Diaminobenzidine using ImmPACT DAB Plus Substrate Kit, Peroxidase (Vector Laboratories) for 1-10 min at RT, followed by hematoxylin staining.

### Statistical analysis

Each treatment was duplicated, and the experiments were repeated at least once to ensure reproducibility. Power analysis was performed to determine the sample size to ensure biological significance. The data were analyzed using FlowJo and GraphPad Prism software. Unpaired student T-tests (two-tailed) were used to analyze the differences between treated and control groups. All statistical data are represented as mean ± SEM. Statistical significance was defined as *P≤0.05, **P≤0.01, and ***P≤0.001.

## Author contributions

RS, ALM, AC, BH, and GY contributed to the conception and design of the study. RS, ALM, AC, FY, DA, SP, HJ, CT, MD, YP, and HZ performed experiments. RS, ALM, AC, BH, and GY performed data analyses. AC, MD, YP, and HZ edited the manuscript. RS, BH, and GY wrote and finalized the manuscript.

## Acknowledgments

The authors would like to thank Dr. Yijen Wu and her team members from Rangos Research Center Animal Imaging Core for assistance with MRI and IVIS imaging; Joshua Michel and his team members from Rangos Research Center Flow Cytometry core for assistance with flow analyses; This work was supported by the NIH UG3TR002842 (GY), the Scientific Program Fund from the Children’s Hospital of Pittsburgh (BH), and the NIH/NCI 1R01CA259124 (BH).

## Figure legends

**Supplementary figure 1. Human cell percentages of the humanized DRAG mice. (A)** Frequency of human CD45^+^ cells in the PBMCs and spleen of DRAG mice. Spleen shows a higher number of human CD45^+^ cells compared with blood. The data for spleen and blood represents 4 and 11 mice, respectively. Accumulation of human CD45^+^ cells in the spleen shows the importance of this secondary lymph organ in the local and systemic regulation of immune cells. **(B-H)** Human stem cells differentiated into CD45^+^ (**B**), CD14^+^ (**C**), CD3^+^ (**D**), CD11b^+^ and CD11c^+^ (**E**), CD4^+^ and CD8^+^ (**F**), CD19^+^ (**G**) and CD56^+^ cells (**H**) with different frequencies in blood PBMCs. Only two mice per marker are shown.

**Supplementary figure 2. Differentiation of human immune cells in the blood and spleens.** Human stem cells differentiated into CD45^+^ (**A**), CD14^+^ (**B**), CD3^+^ (**C**), CD11b^+^ and CD11c^+^ (**D**), CD4^+^ and CD8^+^ (**E**), CD19^+^ (**F**), and CD56^+^ cells (**G**) with different frequencies in spleens. Only two mice per marker are shown.

**Supplementary figure 3. Gating strategies. (A)** Gating strategy for analysis of regulatory T cells (middle panel, including CD25^+^Foxp3^+^CD4^+^ T cells and CD25^+^Foxp3^+^CD8^+^ T cells), naïve and memory CD4^+^ and CD8^+^ T cells (bottom panel, including Naïve CD4^+^ and CD8^+^ T cells (CD45RA^+^CD62L^+^), central memory CD4^+^ and CD8^+^ T cells (CD45RA^−^CD62L^+^), effector memory CD4^+^ and CD8^+^ T cells (CD45RA^−^CD62L^−^) and terminal memory CD4^+^ and CD8^+^ T cells (CD45RA^+^CD62L^−^). **(B)** Gating strategy for analysis of myeloid cells, including monocytes (CD14^+^), macrophages (CD11b^+^), and dendritic cells (CD11c^+^). **(C)** Gating strategy for analysis of various lymphocytic populations, including CD45^+^ (total lymphocytes), CD3^+^ (total T cells), CD4^+^ and CD8^+^ T cells, B cells (CD19^+^), and natural killer (NK) cells (CD56^+^).

