## Supplementary figures and images for "Development of a human glioblastoma model using humanized DRAG mice for immunotherapy"

### Supplementary Figures

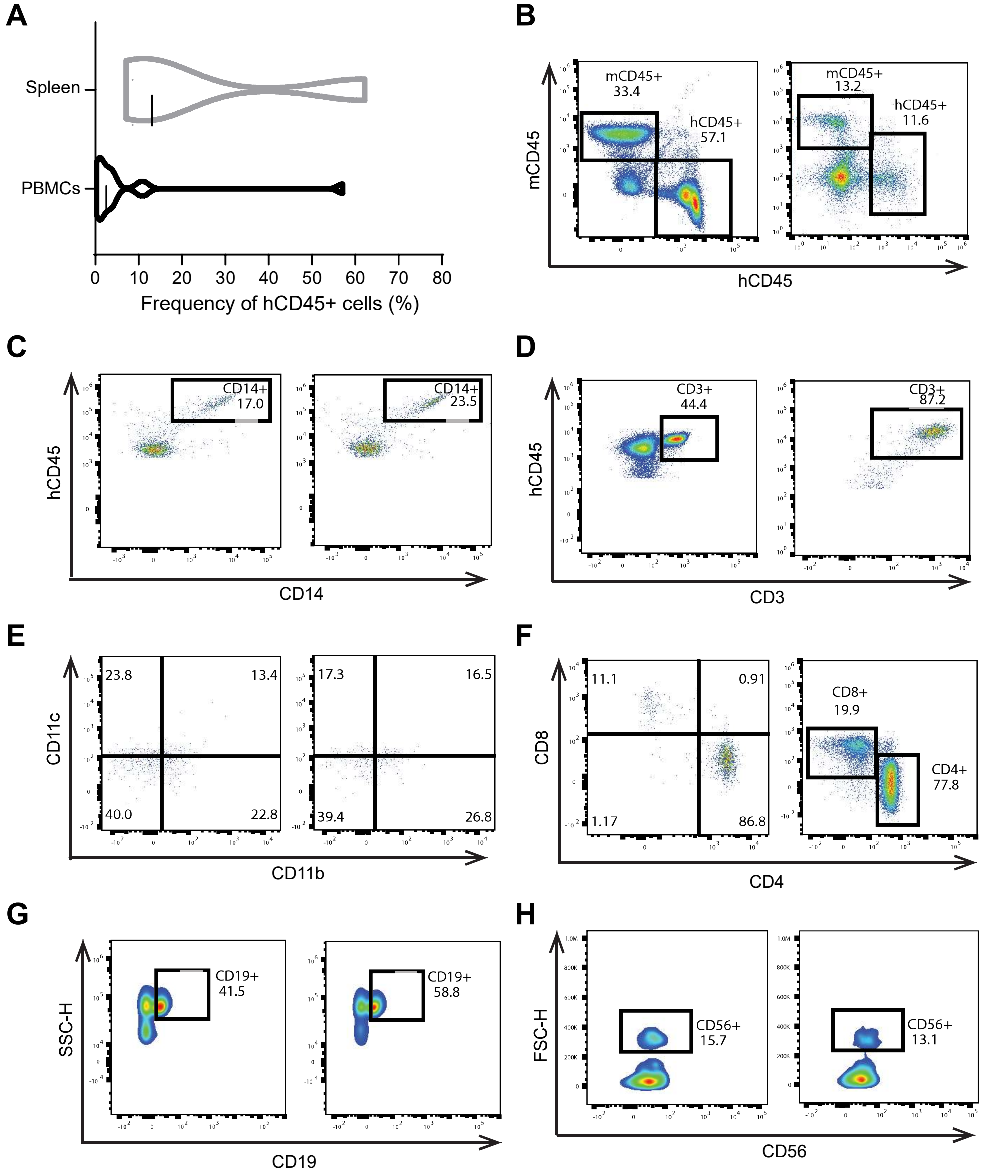

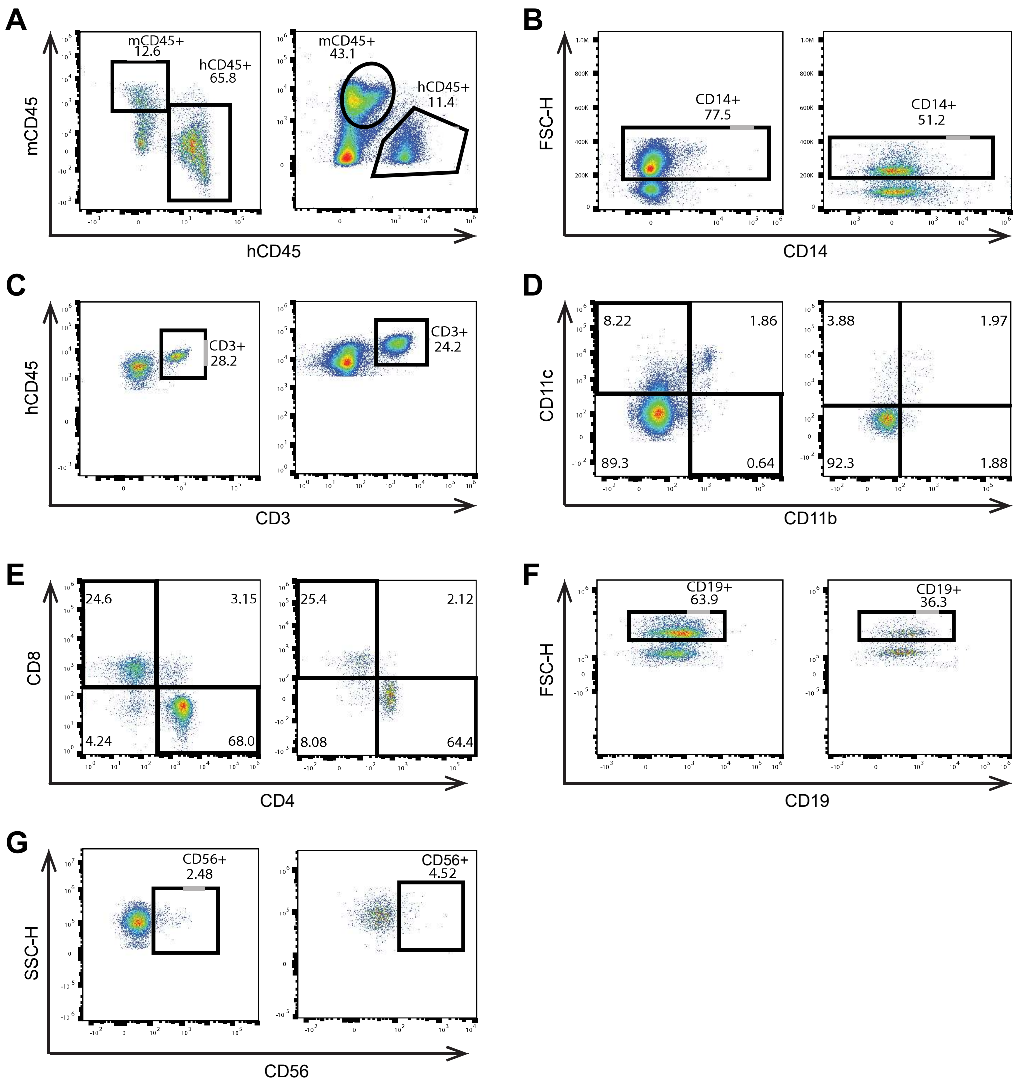

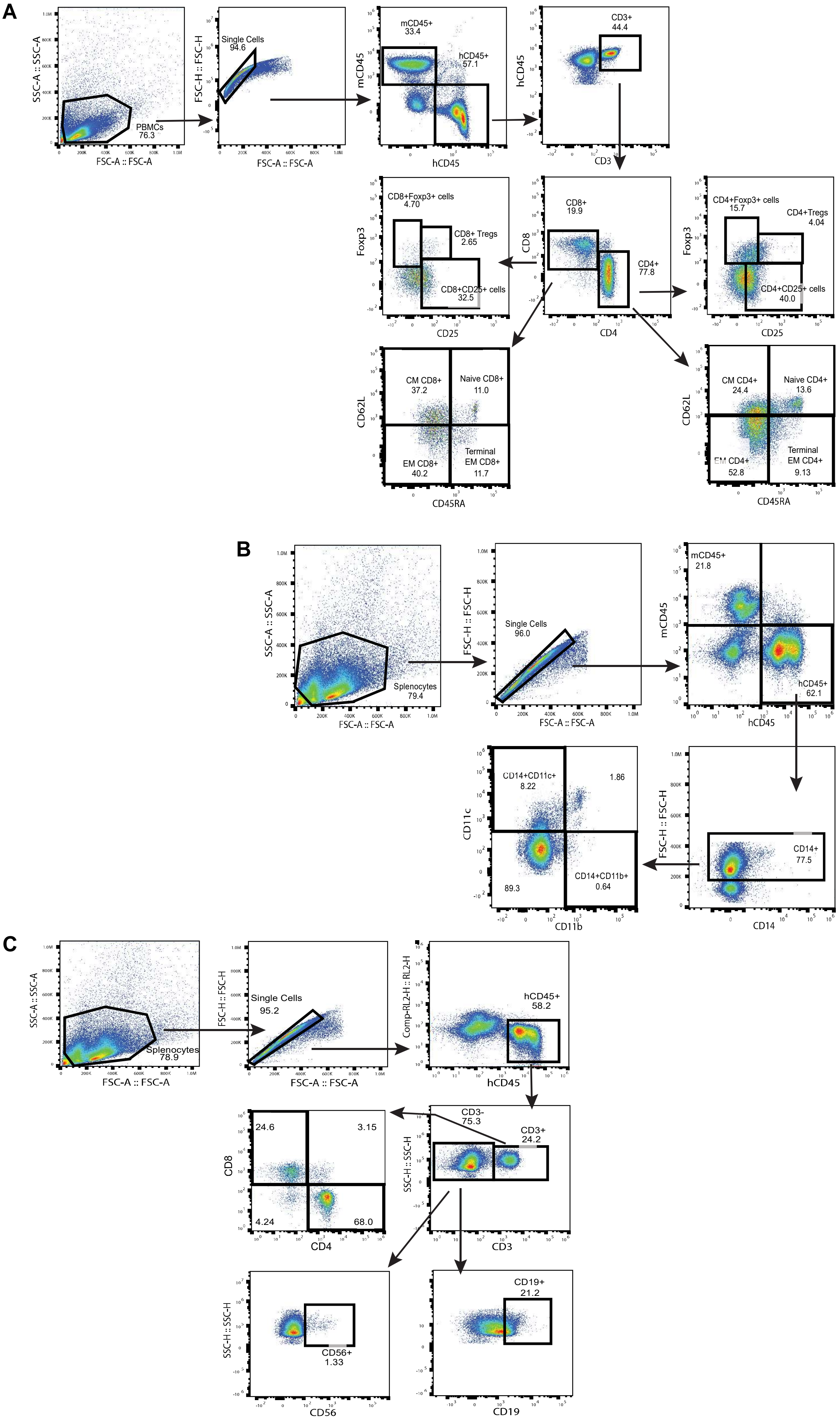
